# 5’-NADylation of RNA is well-tolerated by bacterial cell

**DOI:** 10.1101/2023.03.31.535138

**Authors:** Jana Wiedermannova, Ravishankar Babu, Yulia Yuzenkova

**Affiliations:** CBCB, FMS, Newcastle University; NUBI, FMS, Newcastle University

## Abstract

Recently discovered ability of various RNA polymerases (RNAPs) of bacteria and eukaryotes to initiate RNA synthesis with cofactors such as NAD^+^/NADH (nicotinamide adenine dinucleotide). This phenomenon was quickly dubbed “non-canonical capping” as the extra 5’ moieties of RNA resemble eukaryotic m^7^G cap superficially. NAD must outcompete abundant ATP, to serve as initiating nucleotide. It is still unclear which physiological conditions permit significant RNA NADylation and to what degree NADylation serves typical functions of a canonical cap in bacteria.

Here we found that the extent of NADyation depends primarily on fidelity of transcription initiation –NAD concentration, specific contacts of RNAP active site with NAD/ATP alternative substrates, and genomic DNA supercoiling. It is much less affected by posttranscriptional processes such as de-capping, processing by main 5’-dependent ribonucleases RNaseE and oligoribonuclease. 5’-NAD inhibits neither translation of a leaderless 5’-NAD-RNA nor a physiological base-pairing of small regulatory NADylated RNA with its antisense target RNA. Translation exposes 5’-NAD for de-capping by NudC enzyme.

## Introduction

NAD and other cofactors and small molecules were found as 5’-terminal modification of RNA in bacteria, eukaryotes and viruses (1–3). So far analogues of ADP, GDP and UDP were identified at 5’-end of RNA. 5’ RNA NADylation is one of the most abundant and better-studied modification. Various unrelated RNA polymerases install NAD on RNA using it as non-canonical initiation substrate for RNA synthesis (4, 5). The 5’-NAD has a superficial resemblance to the typical m^7^G cap found in eukaryotes, which is why it was given a name of a non-canonical cap (NCC). Only a small portion of a given RNA is NADylated, but up 70% of some RNA species, e.g. sibD of *E. coli* were reported NADylated (6). By analogy with canonical m^7^G capping, de-capping (deNADing) enzymes were proposed to exist in both Kingdoms, in *E. coli* this function is played by NudC, a NUDIX hydrolase (enzymes that catalyse the hydrolysis of nucleoside diphosphates linked to other moieties X) (7). Many fundamental questions about NADylation remain unanswered, especially in bacteria, where NADylation is arguably most important in the absence of canonical caps. The main problem is whether NADylation in bacteria plays roles similar to those of classic m^7^G eukaryotic cap – RNA fate/function and translation efficiency. Other questions are how NAD incorporation is regulated and whether it is connected to physiology of the cell.

It is still unknown how the 5’-NAD affects processing of RNA in *E. coli*. So far only RNAI, small RNA regulating replication of plasmids with ColE1 origin was shown stabilised by NAD cap in *E. coli* in the absence of de-capping enzyme NudC (5). Would cap present any obstacle for RNA processing by nucleases? Main RNA processing enzyme of *E. coli*, RNase E cleaves RNA internally, but its activity can be determined by the nature of 5’ end of RNA. After fragments created by RNase E are degraded by 3’ exonucleases, degradation RNA is completed with Orn, a highly specific dinucleotide hydrolase (8). If Orn tolerates capped dinucleotides (here and further the length of RNA is in conventional nucleotides) is not known. Many highly capped RNA of *E. coli* are regulatory sRNA (1), via base-pairing they regulate their target mRNAs stability, or either repress or stimulate translation.

De-capping nuclease NudC was co-isolated from *E. coli* cells with ppGpp, stress alarmone involved in bacterial adaptation to stationary phase/starvation (9). Potential ppGpp modulation of NudC activity may provide a direct link between capping and physiological conditions/various stresses known to lead to ppGpp accumulation, and to explain why capping is higher in stationary phase (6).

How NAD+ cap affects translation of mRNA in bacteria is largely unknown. For eukaryotes there being conflicting reports – in yeast and human cells capped transcripts are destined for degradation and presumably not translated, yet in plants they were found in polyribosome fraction/translated (10, 11). Bacterial translation machinery does not rely on 5’-end modification. Yet in case of leaderless mRNA non-canonical cap may directly affect assembly of translation initiation complex, since the initiating codon AUG is changed to NADUG. Leaderless mRNAs are found in all domains of life, widespread in bacteria, frequent especially in *Mycobacteria, Streptococcus* and *Deinococcus* phyla.

## Results

### Determinants of NAD incorporation into an RNA – [NAD]/[ATP] ratio, DNA accessibility, supercoiling and coordination of substrate in the RNAP active site

Extent of RNA NADylation potentially depends on two parameters – absolute NAD concentration and its ratio to canonical RNAP substrate, ATP. ATP is present in higher concentration under standard growth conditions (12), directly competes with NAD as initiating substrate; *in vitro* ATP apparent *K_M_* is lower (4) and the specificity constant, V_max_/*K_M_* is favourable (5). How levels of RNA NADylation depend on cellular concentrations of these two major metabolites, and what conditions would favour NAD incorporation?

To answer this and further questions related to physiological context of NADylation of RNA, we measured ATP and NAD concentrations with commercial kits. For quantification of the bulk RNA NADylation, we developed a fluorescent method, FluorCapQ. This sensitive, simple and reproducible method to measure NAD on RNA is a modification of an assay developed by Putt et al. (13). The method is based on properties of N-alkylpyridinium compounds (e.g. nicotinamide moiety of NAD), which convert to fluorescent molecules through reaction with a ketone (e.g. acetophenone) followed by heating in excess acid (14). The resulting fluorescent compound can be quantified (see Materials and Methods for details). The sensitivity and linear range are superior compared to previously reported colorimetric NAD-CapQ method (15). Importantly, the concentrations of NAD-RNA based on FluorCapQ method are close to methods based on LC-MS (7 vs 2.5 fmol NAD/μg RNA, respectively (2).

We found that in most widely used laboratory growth conditions for *E. coli*, i.e. LB medium, aerobic growth with shaking in a flask, intracellular NAD concentration is about ~250 μM. Importantly, this concentration is below ~400 μM *K_M_* for NAD^+^ in transcription initiation on promoter for RNAI gene (4).

To vary NAD concentration, we have employed NAD auxotrophic strain of *E. coli*, YJE004, where level of intracellular NAD can be manipulated by varying concentration of NAD in growth media (16). In YJE004 strain *nadD* gene for NAD biosynthesis is deleted and NAD transporter *Ntt4* from endosymbiont *Protochlamydia amoebophila* UWE25, transporting intact NAD(H) in counter-exchange with ADP is expressed from plasmid vector (16). Using this strain we tested levels of extracellular NAD from 10 μM to 1 mM which translated into NAD concentrations from 20 μM to 570 μM (Fig. 1A). As alternative way to lower down intracellular NAD concentration we used WT strain grown in Gutnick media with limited nitrogen concentration.

**Figure 1.**
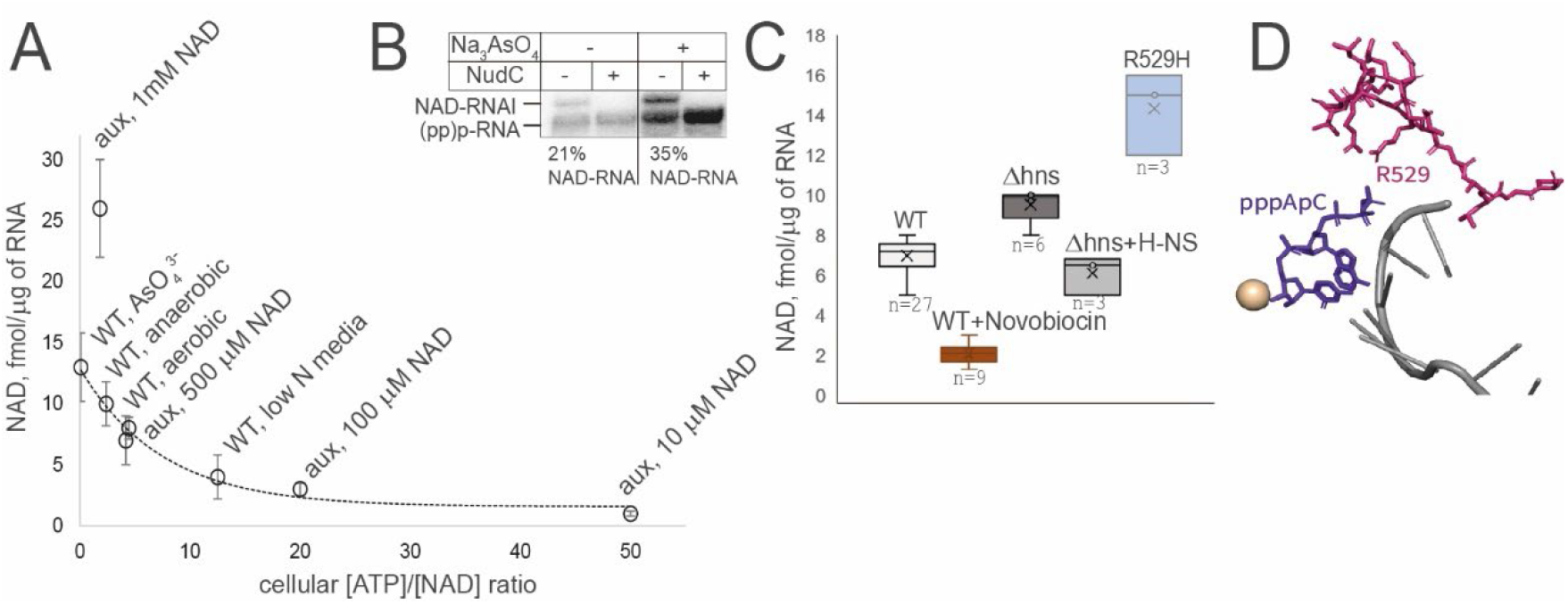
[ATP]/[NAD] ratio, DNA supercoiling and RNAP Rif pocket residue R529 influence cellular RNA NADylation. **A.** RNA NADylation is a function of [ATP]/[NAD] ratio. Plot of NADylated RNA concentration dependency on intracellular [ATP]/[NAD] ratio. **B. Decrease of cellular [ATP] leads to increase of NADylation of RNAI.** Treatment with AsO4^3-^ increased RNAI NADylation. Total RNA was run on boronate gel, treated/untreated with NudC, used for Northern blot with DNA probe against RNAI. **C. Changes in DNA supercoiling, H-NS deletion and RpoB R529H mutation affect NADylation of RNA. D. Structure of RNAP with pppApC RNA.** Mg^2+^ is shown in beige, fragment of β subunit with R529 is in magenta, RNA is in violet.

To test the effect of [ATP], a competing substrate, we inhibited ATP synthesis by placing cells in microaerobic conditions ([ATP] ~ 500 μM) and treated them with AsO4^3-^ (inhibitor of ATP synthase (17)) has led to a significant drop in concentration of ATP to ~30 μM ([NAD] remains unchanged ~220 μM).

In general, steady state levels of NAD-RNA followed the simple trend, correlating with [ATP]/[NAD] ratio, from 1 fmol NAD/1μg of total RNA in YJE004 ([ATP]/[NAD] ≈ 51) to 12.5 fmol NAD/μg of total RNA ([ATP]/[NAD] ≈ 0.09) in cells treated with arsenate.

In parallel, we measured NADylation of RNAI, one of the highest NADylated RNAs of *E. coli* (5), by Northern blots of total RNA samples from untreated cells and those treated with arsenate, run on boronate gels, allowing separation of 5’-NAD and 5’-ppp RNA species (18). Consistently with bulk RNA NADylation measurements, the NAD-RNAI to ATP-RNAI ratio grew from 21% to 35% (Fig. 1B).

The outlier with high level NADylated RNA is YJE004 strain grown with 1 mM NAD supplement, where intracellular level of NAD is ~570 μM. This is the only case where [NAD] rose above its apparent *K_M_* of ~400 μM as the initiation substrate for RNAP.

Nicotinamide and triphosphate moieties of 5’ NAD and 5’ ATP respectively of the short nascent RNA make different contacts with amino acid residues of the RNAP active site the Rifampicin pocket (5), and were shown to influence NAD incorporation *in vitro* (4). We tested mutant strains with substitutions in *rpoB* gene leading to amino acid changes of S512F, Q513L, H526Q, R529H, I572F in FluorCapQ assay. Only R529H mutant led to significant increase in NAD-RNA (Fig. 1C). In the structure R529 coordinates triphosphate of the 5’-ATP, therefore we suggest that R to H change decreased ability to coordinate triphosphate moiety of ATP thus creating preference for NAD incorporation (Fig. 1D).

We found that short treatment of cells with novobiocin, inhibitor of gyrase which leads to decrease in negative DNA supercoiling, lowered down NADylation of RNA. This suggests that genomic DNA supercoiling is one of the major factors controlling incorporation of NAD, and probably promoters prone to NADylation are those sensitive to supercoiling.

To support this hypothesis, we tested levels of RNA NADylation in H-NS deletion strain. H-NS is a major nucleoid organising protein of *E. coli*, condensing DNA into the nucleoid. Deletion of H-NS increases negative supercoiling of chromosomal and plasmid DNA (19). H-NS controls access of the protein machinery required for gene transcription and thereby preventing spurious transcription (20, 21). We found that deletion of H-NS led to increase in overall RNA NADylation, while overexpression of H-NS from plasmid led to decrease in NADylated RNA (Fig. 1C). These results corroborate the results of experiment with novobiocin. Nevertheless, we cannot rule out the alternative conclusion - that at least partially NADylated transcripts originate not from conventional promoters, but are a products of spurious transcription, and removal of H-NS-dependent silencing is known to activate some cryptic initiation sites (21).

### Functional binding of antisense RNAII is not affected by 5’NAD of RNAI

The majority of highly NADylated RNAs in *E. coli* are small regulatory RNAs (5) which function via base-pairing with a target sequence e.g. mRNAs or other sRNAs. This is the case for RNAI, which has its partner RNAII and sib RNAs which base-pair with their corresponding antisence ibs RNAs; gcvB sRNA regulates expression of high number of target genes by base-pairing with their mRNAs (22). RNAII forms a full-length duplex with RNAI, as part of the replication control mechanism of ColE1 origin plasmid. Initially RNAI forms “kissing complex” with RNAII, complex then stabilised by 5’ end of RNAI interaction leading to formation of full length duplex additional base pairs involving the 5’ segment of RNAI (Fig. 2A) (23).

**Figure 2.**
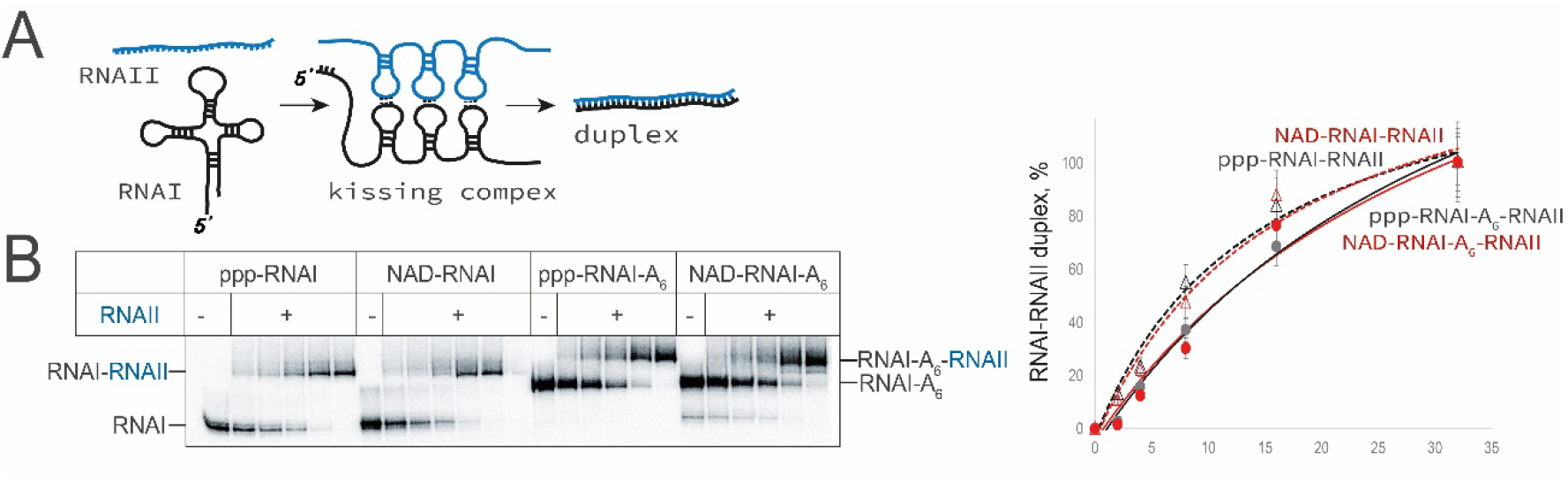
NADylated RNAI forms a duplex with RNAII with same efficiency as triphosphorylated RNAI. **A. Scheme of the RNAI-RNAII duplex formation. B. Duplex formation is not affected by RNAI NADylation.** Labelled RNAI (non-adenylated or hexa-adenylated at3’ end) was incubated with increased concentrations of unlabelled RNAII. Right – the percentage of RNAI in a duplex is plotted against RNAII concentration.

To test if 5’-NAD affects RNAI duplex formation with RNAII, we analysed efficiency of RNA-RNA duplex formation between *in vitro* synthesise RNAI and 128 nt long 5’ end complementary part of RNAII. RNAI was radioactively labelled; RNAI-RNAII duplex migrates slower than single-stranded RNAI under our experimental conditions, and can be observed as a band above RNAI band.

In the experiment on Fig. 2B we have found that affinity to RNAII of ppp-RNAI is the same as that of NAD-RNAI. The main cellular form of RNAI is 3’ hexa-adenylated RNAI. This form has lower affinity to RNAII *in vivo* (24). Indeed, affinities of *in vitro* synthesised adenylated forms of RNAI were lower than non-adenylated, yet ppp-RNAI-A6 and NAD-RNAI-A6 affinities to RNAII were similar (Fig. 2B).

### 5’-NADylated leaderless mRNA is bound by 70S ribosomes and efficiently translated

Classic m^7^G capping is needed for efficient translation of eukaryotic mRNA (25). NAD capping in eukaryotes appears to inhibit translation initiation (26), although this inhibition may be an indirect effect of preferential degradation of the NAD-RNA in the cell. Prokaryotic translation machinery has not evolved an apparent means for specific 5’-end of mRNA recognition. Leaderless mRNA is a special case, where ribosome binds at the very 5’ end. NADylated leaderless mRNA has (NAD^+^)UG start codon instead of AUG. *E. coli* codes for only a few leaderless mRNAs, the best studied model of leaderless translation is mRNA coding for CI repressor protein of bacteiophage λ (27).

Using *in vitro* produced 70-nt long 5’ fragment of CI mRNA with either AUG or (NAD^+^)UG start codon we found that the complexes of these mRNAs with ribosomes and initiating tRNA^fMet^ have the same dissociation constants, ~3.5 nM vs ~4.3 nM, respectively, as measured by Bio-Layer interferometry (Octet bioanalytic instrument) using mRNA immobilised via annealing to biotinylated DNA oligo on solid support. In addition, the position of ribosome and its affinity was tested in a toeprint experiment with RelE toxin, which specifically cleaves mRNA between the second and third nucleotides of the codon in the vacant A-site of the post-translocated ribosome (28) (Fig. 3A). For this experiment we used the same *in vitro* synthesised 70-nt long 5’ fragment of CI mRNA which was labelled at the 3’ end. Ribosome placement was not changed by 5’-NAD, judged by same length of the RelE cleavage fragment. Affinities to mRNAs measured by titration of ribosomes were similar for ppp-RNA and NAD-RNA, 38 nM vs 43 nM, respectively. These values correlated well with *K_d_* values for two mRNAs.

**Figure 3.**
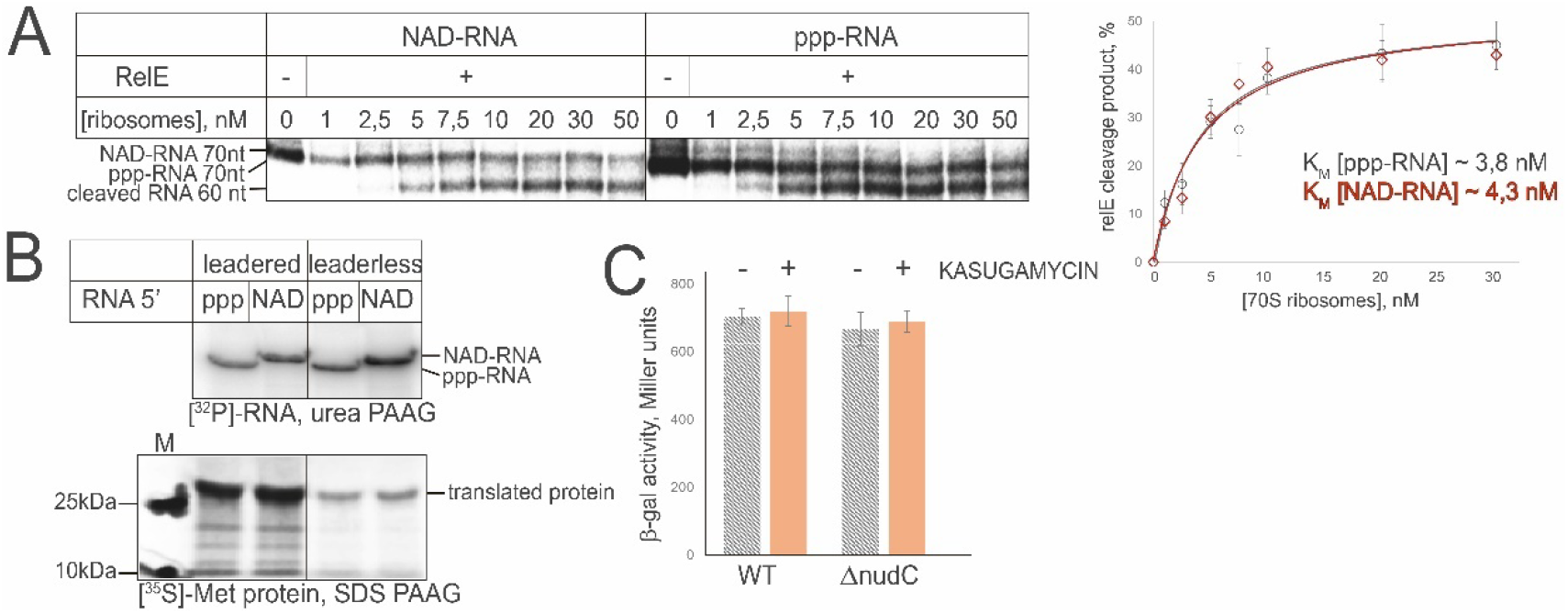
Translation of leaderless mRNA is not affected by 5’-NAD. A. RelE cleavage of NAD- and ppp-RNA at increasing concentration of ribosomes. Left – gel showing positions of original and cleaved RNA, right – plot of percentage of RelE cleaved RNA vs ribosomes concentrations. **B. *In vitro* translation of leadered and leaderless NADylated or triphosphorylated RNA.** Top panel –mRNAs labelled with [α-^32^P] UTP run od denatured urea gel, bottom – corresponding peptide labelled with [^35^S] Met run on SDS gel. **C.** Beta-galactosidase activity of cell lysates expressing CI-lacZ fusion leaderless mRNA construct in either WT or ΔNudC strains with and without Kasugamysin.

To assess the effect of 5’-NAD on the overall efficiency of translation, *in vitro* produced 5’-NAD-mRNA and ppp-mRNA coding for short CI-derived N-terminal 34 amino acid residues-long peptide were used in *in vitro* translation experiment with commercial *E. coli* S30 Extract System with [^35^S]-Methionine. These mRNA produced comparable amounts of 34-amino acid long N-terminal CI-lacZ peptide (Fig. 3B).

To test if NAD-RNA is translated in the cell, we constructed plasmid vector with lacZ gene fused with short initial portion of λ CI gene (10 N-terminal amino acid residues, CI-lacZ), placed under RNAI promoter on plasmid pJW370. This construct, transformed into *E. coli* WT and Δ*nudC* strains, produced comparable β-gal activity (Fig. 3C). To confirm that this signal originates from protein translated from leaderless mRNA, we used Kasugamycin, antibiotic specifically inhibiting leadered but not leaderless translation (29). We observed no significant difference in β-galactosidase activity from any of these strains and conditions. Importantly, as ~ 10% of NAD-RNA is observed on Northern blot (Fig. 4B) it suggests that it is translated, as lacZ mRNA is known for efficient coupling between transcription and translation, and lacZ transcription would have stopped upon translation inhibition (30).

**Figure 4.**
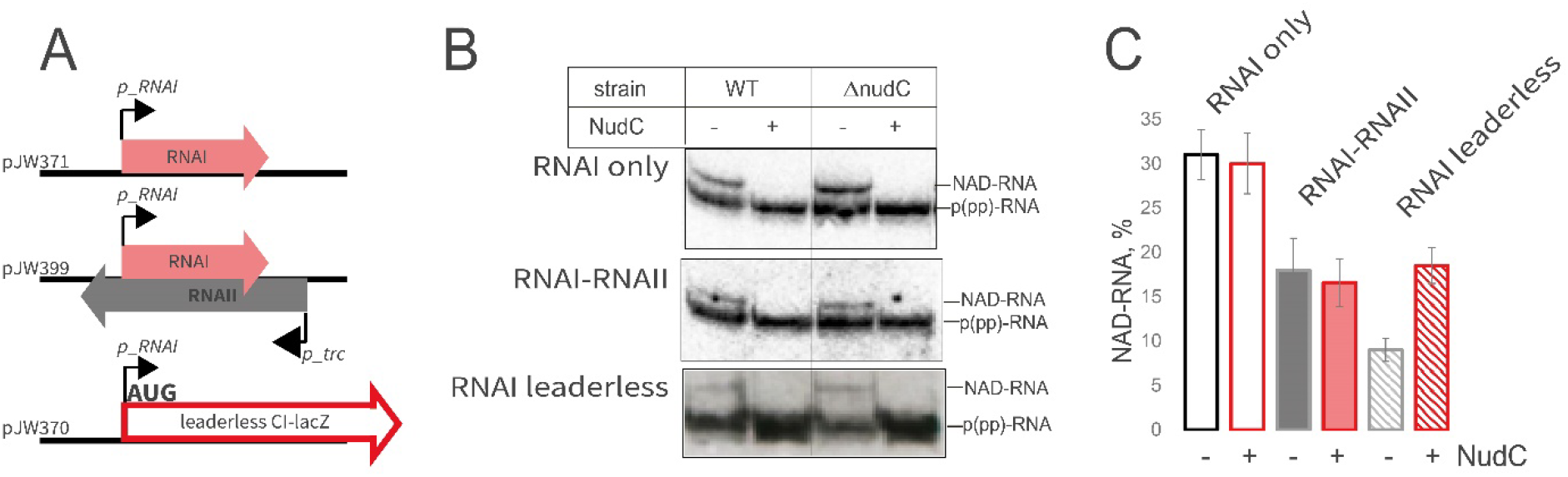
Dependence of RNA NADylation on duplex formation with antisense RNA and translation. **A. Scheme of the plasmids used for experiment. B. Northern blot against RNAi and its derivative.** Total RNA run on boronate PAAG and treated/untreated with NudC, for translated (leaderless) construct total RNA was first treated with DNazyme to produce shorter 5’ fragment. **C. Plot of the ratio of NADylated RNAI produced in these strains.**

### RNA-RNA duplex formation does not protect NAD-RNA, translation exposes 5’-NAD for de-capping

Since 5’-NAD of RNA affects neither duplex formation with antisense RNA nor translation, we wondered whether fate of NADylated RNA is influenced by antisence-RNA or ribosome binding, i.e. whether 5’-NAD is protected, or *visa versa*, exposed. We constructed plasmid vectors pACYC184-based (p15A origin of replication) constructs with either RNAI (from pCDF plasmid with colE1 origin) cloned under its own promoter, or both RNAI and 5’ fragment of RNAII (under strong constitutive *p_trc* promoter and strong terminator upstream of p_RNAI, Fig. 4A). Neither of these extra RNAs interfere with plasmid replication. As can be seen from Fig. 4B, formation of duplex with RNAII slightly lowers down the proportion of NADylated RNAI, compared to plasmid with RNAI only (Fig. 4C). Notably, this proportion remains the same for Δ*nudC* strain, suggesting that formation of RNA-RNA duplex does not affect processing of the RNAI by de-capping enzyme and suggesting that the role of NudC is limited to unstructured RNAs.

Translation, on the other hand, lowered down the proportion of NADylated RNA, and deletion of NudC has a significant effect on it (9% vs 16.5%). This may suggest that the absence of secondary structure and translation process expose 5’-end to NudC de-capping, and ribosome binding is probably too transient to protect the “cap.”

### Processing of NADylated RNA– de-capping and major RNases processing

One of the main functions of m^7^G cap is regulation of mRNA stability (31). This function was proposed for NAD-cap, but opposite effects for eukaryotes and bacteria were observed – while in eukaryotes it speeds up degradation, in bacteria 5’-NAD was suggested to protect RNA, based on the experiment where half-life of RNAI was increased in *ΔnudC* strain (5).

The RNA degradation cascade in *E. coli* is initiated by RNaseE and finalised by oligoribonuclease Orn. Only these two essential enzymes may depend on the 5’ end, and there are no 5’-dependent exonucleases in *E. coli*. We observed the lower proportion of bulk NAD-RNA measured by FluorCapQ in WT strain and in a *E. coli* strain N3431 rne-3071(ts) with temperature-sensitive RNaseE at non-permissive conditions (Fig. 5A). The lower proportion of NAD-RNA would be expected if it is ppp-RNA that gets preferentially degraded in RNaseE depleted cells, suggesting that the degradation pathway based on RNase E is not a major pathway for bulk NAD-RNA.

**Figure 5.**
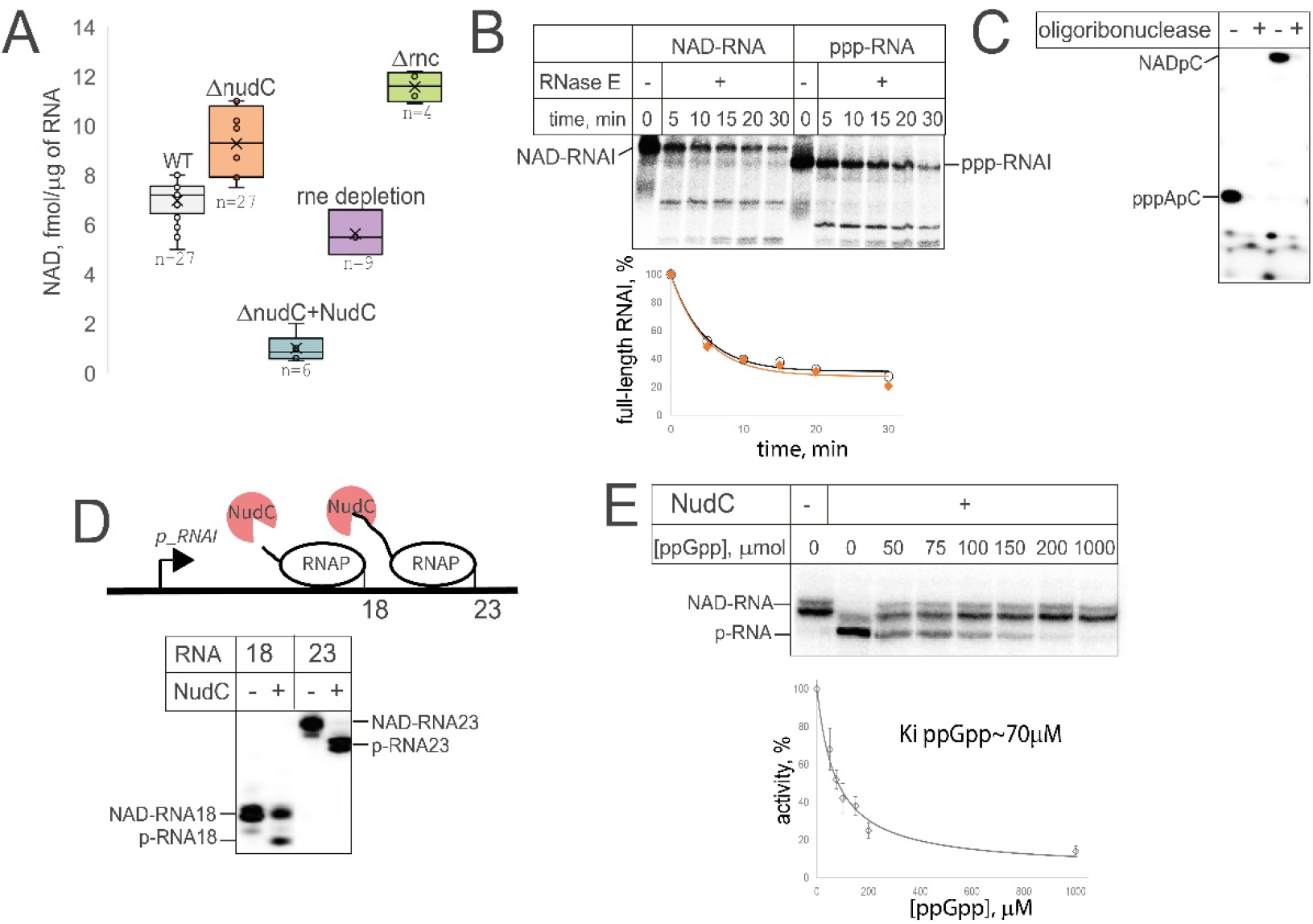
NADylated RNA processing. **A. Amount of NADylated RNA in cells with deletions and depletions of major RNases and decapping enzyme NudC.** Concentration of NAD on RNA was measured by FluorCapQ, Δrnc is a deletion of RNaseIII, rne depletion is depletion of RNaseE at non-permissive temperature in strain with thermosensitive RNaseE, ΔNudC+NudC is a strain with deletion of NudC expressing NudC from a plasmid. **B. Kinetics of *in vitro* NAD- or ppp- RNA cleavage with RNaseIII. C. Hydrolysis of dinucleotide NAD- and ppp- RNA with Orn oligoribonuclease. D. Co-transcriptional RNA decapping by NudC.** Scheme of stopped elongation complexes started from p_RNAI promoter with short nascent RNA emerging from RNA exit channel of RNAP. RNAP stopped at position 18 and 23 by walking along the template *in vitro* using a subset of NTPs. To the stopped elongation complexes NudC was added. **E. Inhibition of NudC with ppGpp.** NAD-RNA was incubated with NudC at indicated concentrations of ppGpp. Below the gel is a plot of NudC activity vs [ ppGpp].

In contrast, RNaseIII deletion caused accumulation of NAD-RNA. Together these results suggest that NAD-RNA is preferentially degraded by RNaseIII, endonuclease acting on RNA duplexes, in agreement with the estimation that most NAD-RNA species are structured sRNA, of which many are shown to duplex with antisense RNAs as part of their regulatory mechanism (1).

RNaseE degradation mechanism is 5’-end-dependent, and it is well-documented that for RNAI the first degradation step is 5’-end proximal cleavage by RNaseE (32). Therefore, NAD at 5’ end of RNA could affect the efficiency of processing by RNaseE. Consistently with the *in vivo* results, *in vitro* NAD-RNA and ppp-RNA RNAI degradation kinetics by RNaseE are the same (Fig. 5B).

The last step of RNA degradation is performed by Orn, the only enzyme able to hydrolyse RNA shorter than five nucleotides to mononucleotides. Would it have any trouble degrading 5’ NADylated dinucleotides? Using dinucleotides produced by *E. coli* RNAP *in vitro* abortive synthesis on RNAI promoter fragment, we found that efficiency of degradation of short NADpC dinucleotide by Orn was the same as pppApC dinucleotide (Fig. 5C).

We found that NudC has only modest effect on NADylated RNA levels, judging from bulk FluorCapQ assay as well as RNAI by Northern blotting (Fig. 1B). However, overexpression of NudC from plasmid vector led to nearly complete removal of NAD-RNA (Fig. 5A). Solving this apparent conundrum, we found that *in vitro*, NudC is de-capping RNA co-transcriptionally as it emerges from RNAP. Nascent RNA becomes susceptible to NudC as soon as it reaches 18 nucleotides, i. e. when only three 5’ nucleotides are exposed (Fig. 5D). These results corroborate our suggestion that the role of NudC is limited to unstructured RNAs, and RNaseIII may play major role in degradation of NADylated RNA in an RNA-RNA duplexes. This result suggests that when the intracellular concentration of NudC is high it can act co-transcriptionally.

In a few publications, it has been noted that levels of NADylated RNA are somewhat higher in stationary phase (6, 33). There are reports from high-throughput proteomics that NudC may bind ppGpp (9). Indeed *in vitro* we found that NudC is inhibited by ppGpp with inhibition constant in the region of ~70 μM, which is in physiological range of ppGpp concentration (~100 μM in stationary phase, and briefly higher during the transition from exponential to stationary(34)) (Fig. 5E).

## Discussion

Here we have tried to put NADylation of RNA into the physiological context of the cell, and showed that NADylated RNA level while depending on many factors, chiefly follows an absolute NAD concentration, which under most conditions remains below *K_M_* for NAD as transcription initiation substrate. NADylation remains a minor modification (reaching 33% of the RNAI transcript) under extreme conditions of low concentration of competing substrate, ATP (which mimics membrane depolarisation/ proton gradient dissipation), or by artificially feeding NAD via orthologous pump - conditions bacteria rarely encounter.

Importantly, here the percentages of 5’-NADylated species for specific RNAs were calculated based on Northern blotting, a direct method. For these reasons it is perhaps unlikely to come across physiological conditions at which a given transcript could be fully NADylated, at least in *E. coli*.

The sensitivity of RNA NADylation to DNA supercoiling may explain why one of the highest NADylated species, RNAI originates from plasmid, where negative supercoiling is stronger. Supercoiling may contribute to the higher NADylation seen in stationary phase (6, 35), along with ppGpp inhibition of major de-capping enzyme NudC. It also suggests that the relatively weak consensus motif of promoters where RNAP is most prone to incorporation of NAD (33), may reflects not sequence requirement, but propensity to assume favourable DNA conformation.

Another conclusion we reached here is that NADylation of RNA is mainly controlled at incorporation step or co-transcriptionally. Very soon after incorporation, the majority of NAD-RNA becomes nearly invisible for cellular machinery. We may consider several reasons behind it. Firstly, most NADylated RNA are structured sRNA whose regulatory mechanism requires its annealing with anti-sense target. Base-pairing kinetics and affinity of NADylated RNA to its partner are not affected, as we showed for RNAI-RNAII pair. Once 5’-NAD-RNA forms a duplex whether in cis or in trans, it becomes resistant to processing by NudC, enzyme mainly active on exposed 5’-NAD. Processing of double-stranded RNA is proceeds mainly via RNaseIII, an endonuclease insensitive to an RNA 5’-end. Secondly, we suggest that concentration of NudC is relatively low in cells, as deletion of NudC has very modest effect on steady-state NADylation levels. Only overexpression of NudC nearly completely abolishes RNA NADylation. Only certain RNAs are susceptible to NudC post-transcriptionally – e.g. those whose 5’ is exposed, e.g. during (leaderless) translation, explaining modest NudC effect. Upon over-expression from multi-copy plasmid, NudC may de-cap NAD-RNA as it is being made, the mechanism we demonstrated *in vitro* even for a very short RNAs with just three 5’ terminal nucleotides of nascent RNA emerged from RNAP.

We found that leaderless NAD-RNA is translated, i.e. ribosome is not inhibited by non-canonical NADUG start codon. *E. coli* codes for only a few leaderless mRNAs, mostly originating from prophages. However, there are species with high proportion of leaderless transcripts, including *Thermus/Deinococcus* (up to 40%) archaeal (~30%) and *Mycobacteria* (~15%) clades (36), and here the effect of NADylation on translation may be more significant. It also appears that 5’-NAD on RNA is not protected by formation of RNA-RNA hybrid or by translation. The process of translation exposes 5’-NAD moiety for RNA de-capping and subsequent degradation.

In other words, pathways for NAD-sRNA and NAD-mRNA processing may differ. NAD-sRNA processed mainly with RNaseIII, either in duplex or on its own. Pathway including RNase E and de-capping by NudC is perhaps more characteristic for non-structured NAD-mRNA.

Overall, we conclude that NADylation is a stochastic RNA modification based on low fidelity of RNAP at the initiation stage. This modification is silent, i.e. goes largely unnoticed by prokaryotic RNA processing and translation machinery. Its ability to lead a quiet parallel life is however remarkable and may result in potential bistable effects. Future research may address the role of NAD-RNA in other species with high intracellular NAD levels and widespread leaderless translation.

## Acknowledgements

This work was supported by Leverhulme Trust Research Grant RPG-2018-437 to YY. We thank Dr Zongbao Zhao, Dalian National Laboratory for Clean Energy, China for YJE004 strain, Prof Ben Luisi, Cambridge University for rne-3071 *E. coli* strain and RNaseE protein.

## Methods

### Strains and plasmids

YJE004 (BW25113/pET15K-NTT4/nadD::cat) NAD auxotrophic strain was a gift from Dr Zhao, Dalian National Laboratory for Clean Energy, China, Δhns, ΔnudC were from Keio collection, Δrnc strain AB301-105 from Yale *E. coli* stock centre, rpoB mutants S512F, Q513L, H526Q, R529H, I572F were spontaneous rifampicin resistant derivatives of WT strain, rne temperature-sensitive strain (Hfr(PO1), lacZ43(Fs), λ-, rne-3071(ts), relA1, spoT1, thiE1) was a gift from Prof Luisi, University of Cambridge. Plasmids pET28*orn* (expressing *E. coli* Orn under IPTG inducible T7 promoter), pJW370 (pACYC184: *p_RNAI* CI (1-30) lacZ), pJW371 (pACYC184: *p_RNAI* RNAI), pJW399 (pACYC184: *p_RNAI* RNAI, antisense *p_trc* RNAII(1-128) T7 terminator) were constructed in this work.

### Proteins

NudC and ORN oligoribonuclease were overexpressed and isolated as in (4). RNase E protein was a gift from Prof Luisi.

### Bacterial growth

*Escherichia coli* (*E. coli*) strain K-12 BW25113 was used as a wild type strain in this study. Cultures were grown at 37°C with shaking (200 rpm) in Luria-Bertani (LB) broth in aerobic conditions. For anaerobic growth, cultures were grown in a 60 ml syringe containing several sterile glass beads for more efficient shaking, with the same media and shaking conditions as aerobic growth. Novobiocin (Sigma-Aldrich; 25 μg/mL) was used to treat *E. coli* cultures for 5 minutes at 37°C with shaking before being harvested for analysis. NAD auxotrophic strain was grown in LB supplemented with 50 μM NAD overnight and then diluted 100 times into LB with the required concentration of NAD (1uM - 1mM). Cells were harvested for RNA isolation and NAD/ATP quantification during the early stationary phase (OD600,~ 1.0), unless overwise indicated. To limit nitrogen levels, cells were grown in Gutnick minimal media supplemented with 0.2% glucose and nitrogen source. Inhibition of proton pump by AsO4^2-^ was done in LB medium supplemented with 10 mM sodium arsenate (NaAsO4) at OD600 =1 for 15 minutes. Cultures were harvested by centrifugation at 8000g for 2 min to remove medium. The cell pellet was flash frozen in liquid nitrogen and stored in −80°C until used.

### Determination of NAD(H) and ATP levels in *E. coli* cells

100 ul of cell culture were harvested by centrifugation at 9000 g for 1 min at room temperature, the supernatant was decanted, and the cell pellet was flash frozen in liquid nitrogen and stored in −80°C until used. Total cellular NAD levels were quantified using the Sigma-Aldrich NAD/NADH Quantification Kit (MAK037). Free ATP quantification was quantified with ATP bioluminescence assay kit, (Roche, cat.NO.11699709001), according to manufacturers’ instruction.

### RNA isolation and purification

Total RNA was extracted from *E. coli* cells grown at 37 °C in LB medium until the OD600 specified in experiments. For strains containing plasmid pCA NudC, expression of NudC was induced with 0.5 mM isopropyl-β-D-thiogalactoside (IPTG) at OD600 = 0.8. Cells were harvested by centrifugation at 9000 g for 1 min at room temperature, the supernatant was decanted. The cell pellet was then flash frozen in liquid nitrogen and stored in −80°C until used. For RNA extraction, the frozen cell pellet was resuspended in 1 ml of TRI reagent (Sigma Aldrich; T9424) per 3 OD units of bacterial culture (equal to number of cells in 3 ml of a culture of OD_600_ =1) and lysed by incubation at 70°C for 10 min, following the manufacturer’s instructions. The sample was then cooled on ice, and 200 ul of chloroform was added. The mixture was vortexed vigorously for 30 s and then centrifuged at 4°C, 20,000 g for 10 min. The upper phase was extracted twice with the same volume of phenol/chloroform solution (5:1 ratio), and the RNA was precipitated with 2.5 M ammonium acetate and 3 volumes of ethanol in −20°C for 1 hour. The RNA pellet was collected by centrifugation at 4°C, 20 000 g for 30 min, washed with 70% cold ethanol, air dried for 10 minutes, resuspended in 50 μl of RNAse-free water and stored in −80°C until use. RNA quality and concentration were determined using a NanoDrop spectrophotometer.

### FluorCapQ quantification of NAD capping on total RNA

RNA for FluorCapQ was further purified to remove free NAD that would interfere with the analysis and bias the results. RNA was dissolved in 2 M urea and 10 mM Tris pH 7.6, heated for 2 minutes at 65°C, and purified using the Monarch^®^ RNA Cleanup Kit (50 μg) (New England Biolabs) according to the adjusted manufacturer’s instructions for the purification of RNA ≥ 15 nt. The RNA yield was quantified using a NanoDrop spectrophotometer. 10 μg of RNA was dissolved in 40 μl of RNase-free water and mixed with 15 μl of a 25 mM solution of 2-acetyl benzofuran (dissolved in ethanol) and 15 μl of 0.5 M KOH. The mixture was incubated on ice for 20 minutes, then 70 μl of formic acid was added and the mixture was incubated at room temperature for an additional 20 minutes. Standard curve samples were prepared in the same way, just dilutions of NAD were used instead of total RNA. The resulting solution was transferred to a black 96-well microplate (Fluotrac, Greiner) and fluorescence was measured by BMG Clariostar fluorimeter (BMG Latec) (using excitation wavelength of 420±40 nm, emission wavelength of 480±40 nm). The NAD levels in each RNA sample were calculated by comparing the absorbance values to a standard curve generated using known concentrations of NAD. Controls: To ensure the accuracy and reproducibility of the results, we included several controls in the NAD quantification protocol, including no RNA controls, no 2-acetyl benzofuran controls, and known NAD standards.

### Northern blotting and boronate acrylamide electrophoresis for detection and quantitation of NAD-capped RNA *in vivo*

RNA samples were treated/untreated with 400 nM NudC in NudC buffer as specified, precipitated, resuspended in RNA loading dye and denatured (20 mM EDTA, 0.025% bromophenol blue, 0.025% xylene cyanol, 7M Urea, 1x TBE, 100 mg/ml heparin, and 80% formamide at 95°C for 10 minutes). NCIN capping of abundant RNAs was analyzed by a procedure consisting of: (i) electrophoresis on 7.5 M urea, 1x TBE, 10% polyacrylamide gels supplemented with 0.33 % 3-acrylamidophenylboronic acid (APB; Sigma Aldrich); (ii) transfer of nucleic acids to a BrightStar™-Plus Positively Charged Nylon Membrane (Invitrogen) using a semidry transfer apparatus at 15 V for 1 hour (Bio-Rad); (iii) immobilization of nucleic acids by UV crosslinking (UV crosslinker (AH) at 1200 μJ/cm2); (iv) prehybridization 30 min in 37°C in SES1 buffer (0.5M sodium phosphate buffer pH7.2, 7% SDS, 10 mM EDTA pH 8), (v) incubation with a 32P-labelled oligodeoxyribonucleotide probe complementary to the target RNAs (^32^P-labelled using T4 polynucleotide kinase and [g ^32^P]-ATP [Perkin Elmer]) overnight in 37°C in SES1 buffer; (vi) washing 2 times 20 minutes with washing buffer (0.1% SDS, 1xSSC (Thermo Scientific); and (vii) storage-phosphor imaging (AmershamTyphoon). Bands corresponding to uncapped and NCIN-capped RNAs were quantified using ImageQuant software. The percentages of NAD+ -capped RNA (5’-NAD) to total RNA were determined from three biological replicates.

### Detection of low abundant RNAs by Northern blotting with RNA probe

Less abundant sRNAs/mRNAs were detected using radiolabelled RNA probes prepared by T7 *in vitro* transcription with a^32^P UTP (see the section “In vitro transcription with T7 RNAP”). Template for sibD probe was amplified from *E. coli* genome using primers JW253: GATCCGAATAATACGACTCACTATTGGAAAGCCCCTCCCGAAGAA and JW254: ACAAGGGTGAGGGAGGATTTCTC. The RNA probe was purified using Micro Bio-Spin™ P-6 Gel Columns (Biorad), denatured 5 min in 95°C and cooled on ice. The membrane with immobilized RNA was prehybridized in buffer (1% SDS, 6X SSPE (Invitrogen), 10X Denhardt’s Solution (Invitrogen),1M NaCl, sodium phosphate buffer pH 7.4, 50% formamide, yeast tRNA 200 mg/ml (Sigma Aldrich)) for 1h in 42°C and hybridized with the RNA probe overnight at 42°C. The membrane was washed two times with 2x SSC for 10 min, 2x with wash buffer (2X SSC, 1% SDS) at 65°C for 30 minutes and two times with 0.1xSSC at 65°C for 30 minutes. The other steps of the procedure were same as described above.

### DNAzyme cleavage of RNA

Cellular RNAs longer than 200 nt (cI-lacZ fusion mRNA) were processed with DNAzyme before the electrophoresis step: 40 mg of total RNA was mixed with 1 mM DNAzyme (JW131) in buffer containing 10 mM Tris pH 8.0, 50 mM NaCl, 2 mM DTT (total volume 50 ml). Samples were heated to 85°C for 5 min and cooled to 37°C (1°C per 30 s). MgCl2 was added to a final concentration of 10 mM and, when present, NudC was added to 400 nM. Reactions were incubated for 60 min at 37°C, then 100 ml of DNAzyme stop solution (0.6 M Tris-HCl pH 8, 18mM EDTA pH 8, 0.1 mg/ml glycogen) and 500 ml of ethanol were added. Samples were centrifuged (30 min, 21000 g, 4°C), the supernatant removed, and the pellet resuspended in RNA loading dye.

### *In vitro* transcription with T7 RNA polymerase: NAD-RNA/ppp-RNA, size standards for Northern blots

Specific RNAs used in this study were generated by in vitro transcription using T7 RNAP. DNA templates containing class I T7 promoter (for transcripts starting with +1G) and class II T7 promoter (transcripts starting with +1A or NAD) were produced by PCR and purified by PCR purification kit (Qiagen) by manufacturer’s instructions.

To generate size standards for Northern blot analysis, we produced RNAs using T7 RNA polymerase with a class II promoter. Two types of RNA were generated: triphosphorylated RNA and NAD-capped RNA. The RNA synthesis reactions were carried out in T7 transcription buffer containing 40mM Tris-HCl (pH 7.9), 10mM MgCl2, and 5mM DTT. For the synthesis of triphosphorylated RNA, the reaction mixture contained 1mM each of ATP, CTP, GTP, and UTP. For the synthesis of NAD-capped RNA, the reaction mixture contained 1mM each of CTP, GTP, and UTP, 0.2 mM ATP, and 4 mM NAD. Usually 6 pmol of T7 RNAP was used in 100 ul reaction volume. After incubating the reactions at 37°C for 2 hours, the RNA was purified using Micro Bio-Spin™ P-6 Gel Columns (Biorad) according to the manufacturer’s instructions. When preparing radiolabelled RNA, UTP was reduced to 0.1 mM and complemented with 1ul of a^32^P UTP (Perkin Elmer). The purified RNA was then analyzed on a denaturing polyacrylamide gel to confirm the expected size of the RNA products.

### RNAI-RNAII binding assay

The assay was done as described before(Tomizawa, 1984; Xu *et al*., 2002)with few adjustments. Briefly: radiolabelled RNAI species (*NAD-RNAI, ppp-RNAI, NAD-RNAI-A_6_, ppp-RNAI-A_6_)* were in vitro transcribed, analysed and quantified by PAA gel electrophoresis and radiography, to use the same amount of each RNA per reaction.

RNAII was DNAsed, purified by Monarch^®^ RNA Cleanup Kit (New England Biolabs) and quantified by nanodrop. All RNAs were denatured in water (85°C for 2 min, cooled on ice). An equal mass of radiolabelled RNAI was mixed with 2x serial dilution of RNAII (32, 16, 8, 4, 2 and 0 ng) in binding buffer (20 mM Tris-HCl pH 7.6, 100 mM NaCl, 10 mM MgCl2). The final volume of the reaction was 5 ml and it was kept at 37°C for 8 min. The reaction was stopped with 10 ml of gel loading buffer (7M urea, 20mM TRIS-HCl pH 7.6, 0.5% SDS, 10 mM EDTA, 0.005% bromophenol blue, 0.01% xylene cyanole) and 3 ml of each sample were loaded without heating on 10 % polyacrylamide gel containing 8M urea in 1x TBE buffer. The electrophoresis was run in a pre-cooled 1xTBE buffer in a cold room at 8°C for 10 min at 400V for 6h.

### Toeprint experiment with RelE toxin / Binding of 70S ribosome to leaderless RNA, RelE assay

Purification of components for ribosome binding to mRNA and RelE cleavage was done as described previously: 70S ribosomes (Castro-Roa & Zenkin, 2015), RelE (Pedersen et al., 2003) (plasmids kindly provided by Kenn Gerdes, Copenhagen University). tRNA^fMet^ was isolated by an adjusted method developed by Yokogawa et al. (37) using a complementary biotinylated probe JW32 (TTATGAGCCCGACGAGCTACCAGGCT-Biotin) and streptavidin agarose (Invitrogen). Briefly: streptavidin agarose was washed 3 times in 500 ml of hybridization buffer (10mM Tris-HCl (pH 7.6), 0.9 M NaCl, 0.1mM EDTA). 200 ml of biotinylated oligonucleotide JW32 (100 mM) was mixed with 100 ml of washed streptavidin agarose and 5 mg of tRNA from E. coli MRE 600 (Roche) in 1 ml of hybridization buffer. The mixture was heated to 70°C for 10 min and cooled slowly to room temperature while agitating. Agarose was washed 4 times with 400 ml of washing buffer (10 mM Tris-HCl, pH 7.6). The captured tRNA^fMet^ was then eluted twice by heating the beads to 70°C for 5 min in 100 ml of wash buffer, fast spinning, and taking the supernatant. RNA was precipitated and resuspended in nuclease-free water.

Model leaderless mRNA (70 nt long 5’ portion of CI protein of phage lambda) containing 5’-ppp or 5’-NAD was synthesized by T7 RNAP using a template containing class II T7 promoter as described in section for *in vitro* transcription with T7 RNA polymerase. The transcription template was amplified using oligo JW100 as a template and JW1, JW2 as PCR primers. mRNA was radiolabelled at the 3’ end using T4 RNA ligase I (NEB) and [^32^P] pCp (Perkin Elmer) according to the manufacturer’s instructions just the reaction was performed for 1 h at room temperature. The RNA was purified by Monarch^®^ RNA Cleanup Kit (New England Biolabs). Binding of ribosomes to leaderless mRNA was tested by assembling 10-500 fmol of 70S ribosomes with 10 fmol of NAD/ ppp-leaderless mRNA in translation buffer (10 mM Tris-HCl pH 7.4, 60 mM NH4Cl, 10 mM Mg(OAc)2, 6mM β-mercaptoethanol) in the presence or absence of 10 pmol of tRNA^fMet^. The mixutere was incubated at 37°C for 10 min. When indicated, 12 pmol of RelE was added and incubated at 37°C for an additional 10 min. The reaction was stopped by adding 10 ml of transcription loading dye and run on 15% polyacrylamide gel, dried, exposed to storage phosphor screens, scanned and visualised by Typhoon, Cytiva and quantified by ImageQuant software.

### Affinity kinetics measured by Octet^®^ RED96e

The components used for RelE toeprinting assay were used for measurements of affinity constants by Octet^®^ RED96e using Bio-Layer Interferometry (BLI) based on fiber-optic biosensors.

200 nM of the model leaderless mRNA (70 nt CI RNA; see Toeprint experiment with RelE toxin) was hybridised to biotinylated DNA oligo (JW10; 100 nM) in annealing buffer (10 mM TRIS-HCl pH 6.8, 50 mM NaCl) by heating to 65°C for 5 min and cooling 1°C per 30 s to room temperature. The streptavidin sensor of Octet^®^ RED96e was equilibrated in an annealing buffer and the RNA-DNA hybrid was loaded on the sensor. The sensor was blocked with biotinylated protein A to prevent unspecific interactions and washed with translation buffer (10 mM Tris-HCl pH 7.4, 60 mM NH4Cl, 10 mM MgOAc). 202 and 36 nM 70S ribosomes were mixed with 2-fold molar excess of tRNA^fMet^ in translation buffer and loaded to the sensor with the attached DNA-RNA hybrid. The binding and dissociation of 70S ribosomes were monitored and affinity constants were calculated.

### β-galactosidase assay of proteins coded by leaderless mRNA

A short initial portion of phage λ CI gene (10 N-terminal amino acid residues) was fused with lacZ gene and cloned into pACY184 under RNAI promoter resulting in plasmid JW370. This construct produced a leaderless mRNA coding CI-lacZ fusion protein and was transformed into E. coli WT and ΔnudC strains. These strains were grown in LB medium until the exponential phase and then treated with x mM kasugamycin (final concentration of 750 μg/ml) for 30 min. β-galactosidase activity of CI/β-gal fusion was measured as described (38).

### Preparation of RNA template for *in vitro* translation

The RNA template for translation (leadered or leaderless phage lambda CI mRNA) was prepared by T7 RNAP dependent *in vitro* transcription using dsDNA as a template. dsDNA templates were amplified by PCR using plasmid JW195 and oligos JW114, JW115 as primers for the leadered template and plasmid JW190 and oligos JW115, JW116 for the leaderless template. The resulting dsDNA products contain class II T7 promoter and 288 nt (96 N-terminal amino acids) of phage lambda cI protein starting with AUG codon (leaderless) or with leader sequence from pCDF expression vector (leadered). The mRNAs were produced in two forms: with 5’ppp and 5’NAD as described in the section for *in vitro* transcription with T7 RNA polymerase. mRNA was treated with DNAse I (New England Biolabs) and purified by Monarch^®^ RNA Cleanup Kit (New England Biolabs).

To check the quality and quantity of the mRNA, it was labelled by [^32^P] pCp (Perkin Elmer), resolved on 5% PAA gel containing 3.3% APB and the respective bands were quantified to assure that the same amount of RNA will be used in *in vitro* translation assay.

### *In vitro* translation

*In vitro* translation experiment was performed using *E. coli* S30 Extract System for Linear Templates (Promega), according to the manufacturer’s instructions for radiolabelled translation. The reaction was supplemented with [^35^S]-Methionine (Perkin Elmer) and 2.4 mg of each mRNA template per sample and run for 2 h at 37°C. The translated protein (2 ml) was mixed with loading buffer, denatured and analysed on NuPAGE™ 4 to 12%, Bis-Tris SDS PAGE gel (Invitrogen). Besides, the Novex™ Sharp Pre-stained Protein Standard (Invitrogen) was run on the gel. After the run, each of the pre-stained bands of the standard was labelled with 0.1 ml of diluted [^35^S]-Methionine by piercing the gel. The gel was dried, exposed to a storage phosphor screen, and scanned by Amersham Typhoon.

### *In vivo* antisense RNA protection of capping

RNAI from plasmid pCDF-1b (colE1 origin of replication) under its native promoter was cloned into the plasmid pACYC184 (p15A origin of replication) using Gibson assembly (New England Biolabs) generating plasmid JW371, this plasmid was transformed into wt and ΔnudC, *E.coli* strains. To express the antisense RNA to RNAI (5’ end of RNAII), we cloned a strong p_trc promoter upstream of the 5’ end of RNAII and T7 terminator (efficiently recognized by *E.coli* RNAP (39)) downstream of RNAII fragment. The whole RNAI and 5’ end of RNAII complementary to RNAI is transcribed from the resulting plasmid (JW399). While RNAII to is not full-length, it does not promote replication of the plasmid and it is not a functional origin of replication (40).

Bacterial strains were grown in LB medium and harvested for RNA extraction at exponential (OD_600_=1) and stationary (OD_600_=3) phases. RNA was isolated as described. NAD capping of RNAI was assessed by Northern blotting of total RNA isolated from these strains and separated in 10 % PAA gel containing 0.33 % APB as described. RNAI was detected by a DNA probe (JW198) labelled with T4 polynucleotide kinase (New England Biolabs).

### Processing of NAD-RNA *in vitro* by RNaseE and olgoribonuclease of *E. coli*

To determine the susceptibility of NAD-capped and triphosphorylated RNA *in vitro*, equal amounts of radiolabelled RNAI were incubated with or without 50 μM RNAse E at 37°C for various time (0, 5, 10, 15, 20, and 30 minutes) in reaction buffer (50 mM Tris-HCl, pH 7.5, 50 mM KCl, 5 mM MgCl2). At each time point, the reaction was stopped by adding an equal volume of transcription loading dye. The samples were denatured (95°C, 2min) and separated by 10% PAA containing 0.33 % APB as described.

Dinucleotide substrates for oligoribonuclease (Orn) were prepared in *in vitro* transcription reaction on linear PCR-generated template containing RNAImod (4) with 500 μM ATP or NAD and 25 μM *[α* ^32^P] CTP (Perkin Elmer) using hexa-histidine tagged *E. coli* RNA polymerase. After incubation in reaction buffer (50 mM Tris-HCl, pH 7.5, 50 mM KCl, 5 mM MgCl_2_) for 15 min at 37°C RNA polymerase was removed by addition of Ni-NTA agarose beads and spinning down the suspension. Supernatant containing dinucleotides (ppp-A-p-C or NAD-p-C) was transferred to new tube where 100 nM Orn was added and incubated for further 10 minutes at 37°C. The reaction was stopped by adding equal volume of transcription loading dye. The samples were denatured (95°C, 2min), separated by 33% PAAG and visualized by radiography.

## References

1. H. Cahova, M. L. Winz, K. Hofer, G. Nubel, A. Jaschke, NAD captureSeq indicates NAD as a bacterial cap for a subset of regulatory RNAs. Nature 519, 374–377 (2015).

2. J. Wang et al., Quantifying the RNA cap epitranscriptome reveals novel caps in cellular and viral RNA. Nucleic Acids Res 47, e130 (2019).

3. W. E. Kowtoniuk, Y. Shen, J. M. Heemstra, I. Agarwal, D. R. Liu, A chemical screen for biological small molecule-RNA conjugates reveals CoA-linked RNA. Proc Natl Acad Sci U S A 106, 7768–7773 (2009).

4. C. Julius, Y. Yuzenkova, Bacterial RNA polymerase caps RNA with various cofactors and cell wall precursors. Nucleic Acids Res 45, 8282–8290 (2017).

5. J. G. Bird et al., The mechanism of RNA 5’ capping with NAD+, NADH and desphospho-CoA. Nature 535, 444–447 (2016).

6. H. Zhang et al., Use of NAD tagSeq II to identify growth phase-dependent alterations in E. coli RNA NAD(+) capping. Proc Natl Acad Sci U S A 118 (2021).

7. K. Hofer et al., Structure and function of the bacterial decapping enzyme NudC. Nat Chem Biol 12, 730–734 (2016).

8. S. K. Kim et al., A dedicated diribonucleotidase resolves a key bottleneck for the terminal step of RNA degradation. Elife 8 (2019).

9. B. Wang et al., Affinity-based capture and identification of protein effectors of the growth regulator ppGpp. Nat Chem Biol 15, 141–150 (2019).

10. Y. Zhang et al., Extensive 5’-surveillance guards against non-canonical NAD-caps of nuclear mRNAs in yeast. Nat Commun 11, 5508 (2020).

11. X. Yu et al., Messenger RNA 5’ NAD(+) Capping Is a Dynamic Regulatory Epitranscriptome Mark That Is Required for Proper Response to Abscisic Acid in Arabidopsis. Dev Cell 56, 125–140 e126 (2021).

12. B. D. Bennett et al., Absolute metabolite concentrations and implied enzyme active site occupancy in Escherichia coli. Nat Chem Biol 5, 593–599 (2009).

13. K. S. Putt, P. J. Hergenrother, An enzymatic assay for poly(ADP-ribose) polymerase-1 (PARP-1) via the chemical quantitation of NAD(+): application to the high-throughput screening of small molecules as potential inhibitors. Anal Biochem 326, 78–86 (2004).

14. T. Moriya, A. Kawamata, Y. Takahashi, Y. Iwabuchi, N. Kanoh, An improved fluorogenic NAD(P)+ detection method using 2-acetylbenzofuran: its origin and application. Chem Commun (Camb) 49, 11500–11502 (2013).

15. E. Grudzien-Nogalska, J. G. Bird, B. E. Nickels, M. Kiledjian, “NAD-capQ” detection and quantitation of NAD caps. RNA 24, 1418–1425 (2018).

16. Y. Zhou et al., Determining the extremes of the cellular NAD(H) level by using an Escherichia coli NAD(+)-auxotrophic mutant. Appl Environ Microbiol 77, 6133–6140 (2011).

17. W. L. Klein, P. D. Boyer, Energization of active transport by Escherichia coli. J Biol Chem 247, 7257–7265 (1972).

18. G. L. Igloi, H. Kossel, Use of boronate-containing gels for electrophoretic analysis of both ends of RNA molecules. Methods Enzymol 155, 433–448 (1987).

19. F. J. Mojica, C. F. Higgins, In vivo supercoiling of plasmid and chromosomal DNA in an Escherichia coli hns mutant. J Bacteriol 179, 3528–3533 (1997).

20. S. T. Arold, P. G. Leonard, G. N. Parkinson, J. E. Ladbury, H-NS forms a superhelical protein scaffold for DNA condensation. Proc Natl Acad Sci U S A 107, 15728–15732 (2010).

21. S. S. Singh et al., Widespread suppression of intragenic transcription initiation by H-NS. Genes Dev 28, 214–219 (2014).

22. M. Miyakoshi et al., Mining RNA-seq data reveals the massive regulon of GcvB small RNA and its physiological significance in maintaining amino acid homeostasis in Escherichia coli. Mol Microbiol 117, 160–178 (2022).

23. C. Malmgren, E. G. Wagner, C. Ehresmann, B. Ehresmann, P. Romby, Antisense RNA control of plasmid R1 replication. The dominant product of the antisense rna-mrna binding is not a full RNA duplex. J Biol Chem 272, 12508–12512 (1997).

24. F. F. Xu, C. Gaggero, S. N. Cohen, Polyadenylation can regulate ColE1 type plasmid copy number independently of any effect on RNAI decay by decreasing the interaction of antisense RNAI with its RNAII target. Plasmid 48, 49–58 (2002).

25. Y. Furuichi, A. J. Shatkin, 5’-termini of reovirus mRNA: ability of viral cores to form caps post-transcriptionally. Virology 77, 566–578 (1977).

26. X. Jiao et al., 5’ End Nicotinamide Adenine Dinucleotide Cap in Human Cells Promotes RNA Decay through DXO-Mediated deNADding. Cell 168, 1015–1027 e1010 (2017).

27. A. Resch, K. Tedin, A. Graschopf, E. Haggard-Ljungquist, U. Blasi, Ternary complex formation on leaderless phage mRNA. FEMS Microbiol Rev 17, 151–157 (1995).

28. K. Pedersen et al., The bacterial toxin RelE displays codon-specific cleavage of mRNAs in the ribosomal A site. Cell 112, 131–140 (2003).

29. I. Moll, U. Blasi, Differential inhibition of 30S and 70S translation initiation complexes on leaderless mRNA by kasugamycin. Biochem Biophys Res Commun 297, 1021–1026 (2002).

30. O. Yarchuk, N. Jacques, J. Guillerez, M. Dreyfus, Interdependence of translation, transcription and mRNA degradation in the lacZ gene. J Mol Biol 226, 581–596 (1992).

31. S. Meyer, C. Temme, E. Wahle, Messenger RNA turnover in eukaryotes: pathways and enzymes. Crit Rev Biochem Mol Biol 39, 197–216 (2004).

32. T. Tomcsanyi, D. Apirion, Processing enzyme ribonuclease E specifically cleaves RNA I. An inhibitor of primer formation in plasmid DNA synthesis. J Mol Biol 185, 713–720 (1985).

33. I. O. Vvedenskaya et al., CapZyme-Seq Comprehensively Defines Promoter-Sequence Determinants for RNA 5’ Capping with NAD<sup/>. Mol Cell 70, 553–564 e559 (2018).

34. V. Varik, S. R. A. Oliveira, V. Hauryliuk, T. Tenson, HPLC-based quantification of bacterial housekeeping nucleotides and alarmone messengers ppGpp and pppGpp. Sci Rep 7, 11022 (2017).

35. V. L. Balke, J. D. Gralla, Changes in the linking number of supercoiled DNA accompany growth transitions in Escherichia coli. J Bacteriol 169, 4499–4506 (1987).

36. X. Zheng, G. Q. Hu, Z. S. She, H. Zhu, Leaderless genes in bacteria: clue to the evolution of translation initiation mechanisms in prokaryotes. BMC Genomics 12, 361 (2011).

37. T. Yokogawa, S. Ohno, K. Nishikawa, Incorporation of 3-azidotyrosine into proteins through engineering yeast tyrosyl-tRNA synthetase and its application to site-selective protein modification. Methods Mol Biol 607, 227–242 (2010).

38. S. L. Dove, A. Hochschild, A bacterial two-hybrid system based on transcription activation. Methods Mol Biol 261, 231–246 (2004).

39. J. L. Wiggs, J. W. Bush, M. J. Chamberlin, Utilization of promoter and terminator sites on bacteriophage T7 DNA by RNA polymerases from a variety of bacterial orders. Cell 16, 97–109 (1979).

40. M. Camps, Modulation of ColE1-like plasmid replication for recombinant gene expression. Recent Pat DNA Gene Seq 4, 58–73 (2010).

